# PRESTIGE-ST: Patch Resolution and Encoder STrategies for Inference of Gene Expression from Spatial Transcriptomics

**DOI:** 10.1101/2025.08.11.669796

**Authors:** Maninder Kaur, Amit Kumar, Michele Ceccarelli, Raghvendra Mall, Sukrit Gupta

## Abstract

Spatial Transcriptomics (ST) integrates histology with spatially resolved gene expression, offering rich insights into tissue architecture and function. However, its clinical and large-scale deployment is hindered by high costs, technical complexity, and limited accessibility. To address this, computational pathology methods have emerged to predict gene expression directly from histology images, typically framing the task as a multi-output regression problem mapping image patches to gene expression profiles. While several Convolutional Neural Network (CNN) models have been proposed, little is known about how performance is influenced by (a) the number of trainable parameters and (b) the patch size used for prediction. Moreover, existing studies rely primarily on quantitative metrics and overlook biological relevance of predictions. In this study, we systematically evaluated multiple convolution based models (including a Vision Transformer (ViT) model) with different patch sizes on the Xenium based Autoimmune Machine Learning Challenge (AMLC) dataset. We assessed model performance on both globally expressed genes and subsets enriched for immune or disease associated pathways. Our findings reveal that compact CNNs trained on larger patches outperform deeper models, offering superior accuracy in predicting gene expression, especially for biologically important genes. These insights provide practical guidance for designing efficient and biologically meaningful models in the emerging field of image-based gene expression prediction.

**CCS CONCEPTS:** Computing methodologies → Computer vision; Neural networks; • Applied computing → Computational genomics; Imaging.

## 1 INTRODUCTION

Understanding the molecular landscape of complex diseases *in situ* is critical for advancing precision medicine and targeted therapies. Spatial Transcriptomic (ST) has emerged as a powerful technology that enables the simultaneous measurement of gene expression and tissue morphology within intact histological sections, preserving spatial context [1, 2]. By localizing gene activity within the tissue architecture, ST enhances our understanding of cell–cell interactions, microenvironmental dynamics, and disease-specific tissue organization, with applications across cancer, neurodegenerative, and autoimmune disorders [3].

Despite its transformative potential, the clinical and large-scale adoption of ST remains limited due to several practical challenges [4]. The technology is costly, requires specialized equipment and protocols, and often suffers from limited tissue coverage and throughput. Consequently, there is growing interest in computational approaches that aim to replicate the molecular insights of ST using more accessible imaging modalities, particularly Hematoxylin and Eosin (H&E) stained slides, that are the clinical gold standard for tissue histology. While H&E images offer rich morphological information, they lack direct molecular context, constraining their utility in understanding gene-level mechanisms of disease.

To bridge this gap, numerous studies have proposed Deep Learning (DL) models to predict gene expression directly from H&E images by learning representations that correlate image features with underlying molecular phenotypes [5, 6, 7, 8]. These models typically frame the task as a multi-output regression problem, where an image patch is mapped to a vector of gene expression values. Early models leveraged architectures such as ResNet [9], DenseNet (e.g., ST-Net [10]), and VGG (e.g., DeepSpaCE [11]), while more recent approaches like HisToGene [12] incorporate Vision Transformers (ViTs) to capture long-range dependencies in tissue morphology.

However, several key questions regarding model design choices remain unexplored. First, do shallower convolutional models with fewer parameters perform comparably to deeper or more complex models? Second, how does the size of the input patch—which determines the extent of spatial context captured—affect prediction performance, particularly given the resolution differences across ST platforms (e.g., 50 μm in 10x Visium vs. 5 μm in Xenium)? Third, beyond standard quantitative metrics, how do model predictions hold up in biologically meaningful settings—such as predicting expression of genes with high biological variability, known disease markers, or those involved in immune pathways?

In order to study the above factors, we developed a DL framework to predict 460-dimensional spatial gene expression profiles from H&E stained histology slides as illustrated in Figure 1. We performed a comparative analysis of different models and patch sizes, providing design guidance for deep learning pipelines in computational ST. This work makes the following contributions:

1. Model complexity vs. performance: Comparing architectures of varying sizes (Visual Geometry Group-16 (VGG), ResNet-50 (R50), DenseNet-121 (D121), EfficientNet-B0 (EffNet), and a ViT-based model) to assess whether compact models can match or outperform larger ones.
2. Spatial context: Evaluating the influence of patch size by testing the performances of models trained on image patches of different sizes (128 × 128, 32 × 32, 16 × 16), to understand how local vs. broader spatial context impacts gene expression prediction.
3. Biological relevance: Moving beyond conventional quantitative metrics (Mean Squared Error (MSE), Mean Absolute Error (MAE), Pearson Correlation Coefficient (PCC), and Spearman Rank Correlation Coefficient (SRCC)), we performed model assessment on subsets of genes that are biologically significant, i.e. highly variable genes, and genes enriched in immune and inflammatory pathways.

**Figure 1.**
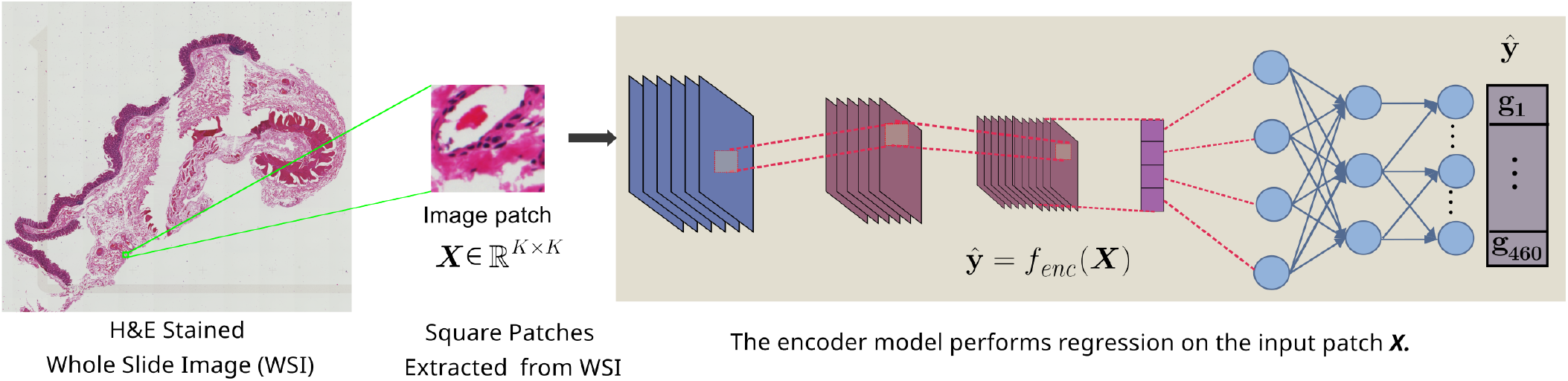
The figure outlines the preprocessing pipeline, where cell-centered H&E image patches are extracted at multiple resolutions (128 × 128, 32 × 32, 16 × 16), and corresponding gene expression profiles are log1p normalized. These patches are then passed to deep learning encoders (VGG, R50, EffNet, D121, and a ViT-based model) to learn feature representations and predict 460-dimensional gene expression vectors, respectively.

Our results on the Xenium ST data from the Autoimmune Machine Learning Challenge (AMLC) [13] show that shallower CNN models trained on larger patches perform comparably or better than deeper CNNs or transformer-based models trained on smaller patches. Notably, the smallest model’s (EffNet) performance in our analysis for biologically relevant genes was the highest for both highly variant and pathway enriching genes. Our results highlight that (i) smaller models (with fewer parameters) can perform as well (and even better) than the bigger models; (ii) neighboring cell information is useful to predict gene expression; and (iii) smaller models are also capable of capturing the inherent biological variance present in gene expression data. These findings offer practical insights for designing effective, lightweight models for computational inference from ST data.

## 2 MATERIALS & METHODS

We model the gene expression profiles of cells from H&E stained Whole Slide Image (WSI) images by using deep learning based frameworks. Below we describe the steps undertaken for data collection, preparation, model building and evaluation.

### 2.1 Dataset Description

This study uses the paired H&E stained images and Xenium ST data from eight colon tissue samples from the AMLC [13]. The dataset includes tissues from six Ulcerative Colitis (UC) patients, the most common subtype of Inflammatory Bowel Disease (IBD), as well as samples from two patients with Diverticulitis (DC), a milder and more localized form of colon inflammation. The UC is further divided into two parts: Inflamed (I) and Non-Inflamed (NI). This dataset, thus, provides a wide range of colon tissue samples, from diseased to healthy. Such heterogeneity can help the model learn and recognize spatial molecular patterns that occur at different stages of health. The included samples comprise:

- DC1, DC5 (diverticulitis)
- UC1_I, UC1_NI (ulcerative colitis, inflamed and non-inflamed)
- UC6_I, UC6_NI (ulcerative colitis, inflamed and non-inflamed)
- UC7_I, UC9_I (ulcerative colitis, inflamed only)

In the base dataset, only partial information was made available for model development, i.e. the ground truth gene expressions were withheld for both the validation and test sets. We therefore created an internal split, where we used cells along vertical strips (total patch count: 55, 743) and random patches (total patch count: 2, 353) from the H&E image as test samples for the gene expression prediction task. Table 1 outlines the distribution of cell-centered image patches used in training and test sets, ensuring sufficient coverage of spatial variability across tissue types.

**Table 1.** Overview of training and custom test data splits. The table reports the number of image patches extracted from each colon tissue sample and their distribution across the training and internal testing groups.

| Sample ID | Total Patches | Training Patches | Test Patches |
| --- | --- | --- | --- |
| DC5 | 140,368 | 132,017 | 8,351 |
| UC1_I | 202,534 | 192,768 | 9,766 |
| UC1_NI | 80,037 | 76,663 | 3,374 |
| UC6_I | 223,790 | 215,396 | 8,394 |
| UC6_NI | 101,485 | 95,714 | 5,771 |
| UC7_I | 144,704 | 133,616 | 11,088 |
| UC9_I | 196,937 | 185,585 | 11,352 |
| **Total** | **1,089,855** | **1,031,759** | **58,096** |

### 2.2 Patch generation from the WSI

H&E stained WSI images from different patient tissue profiles were first processed to generate multiple patches *X* ∈ℝ^*K*×*K*^. These are generated by considering a square of size *K* × *K* around spatial coordinates provided in the corresponding ST dataset. The output *y* ∈ ℝ^*n*^ is a sparse gene expression velctor, which has been log1p normalized (i.e. log 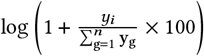 for each patch.

### 2.3 Model training

These generated patches, *X*, are modeled by a set of widely used convolution networks such as R50, D121, EffNet, VGG, and the UNI framework (based on ViT). While, these models can extract meaningful latent feature embeddings from the image patches, we add further Linear layers in order to extract gene expression profiles ŷ ∈ ℝ^*n*^ directly from the model:

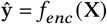

where ŷ corresponds to the predicted gene expressions for the patch, and *f*_*enc*_ represents the encoder backbone followed by an additional regression linear layer. For a regression task, the MSE loss function is used to compute the gradients and update the model weights via backpropagation.

### 2.4 Different Encoder Models

We considered multiple backbone encoders to extract informative features from H&E stained images for gene expression prediction. These models were selected based on their previously shown value in ST studies. They represent diverse architectural paradigms, which allows for evaluating the contribution of both convolutional and transformer-based approaches within the given context.

**VGG** [14], a deep convolutional network, showed strong performance in image-based tasks and has been adapted for a range of medical imaging tasks. Despite its simplicity, VGG has proven to be effective in the field of ST, particularly in DeepSpaCE [11], where it was used to predict gene expression from fixed-size H&E stained image patches. Its layered structure allows for robust feature extraction, making it a natural choice for capturing cellular morphology in our patch-based approach.

**R50** [15] pioneered the concept of skip connections, allowing gradients to propagate more effectively through deeper networks without suffering from vanishing gradients. Also, this architecture has previously been used by Schmauch [9] to predict the bulk RNA-seq data from histopathology patches.ability to capture high-level semantic features makes it suitable for capturing complex tissue structure associated with gene expression.

We also included **D121** [16], which is an upgraded version of ResNet, in which each subsequent layer gets the input from all the previous layers. This dense connectivity ensures better reuse of previously calculated features and mitigates the vanishing gradient problem. DenseNet was previously used in ST-Net [10] for predicting spot-level gene expression from Visium data. This approach highlights its relevance within the spatial transcriptomics domain. The **EffNet** [17] was included in our study due to its strong performance in modeling subtle tissue structure for gene expression prediction. EffNet was adopted in Brst-Net [18], where it was used to predict gene expression from breast cancer histopathology images. Motivated by these findings, we incorporated EffNet as a representative encoder that balances model capacity and computational efficiency.

Finally, we incorporated **UNI** [19] is one of the foundational models for computational pathology, based on large ViT. In order to capture global context and long-range dependencies in high-resolution pathology images, it uses transformer layers. The UNI was trained on over a ≈100 million patches extracted from 100, 000 H&E stained WSI. The dataset contains 20 major tissue types, such as lung, colon, breast, pancreas, *etc*. So far, this is the biggest image encoder for histology. Its inclusion allows us to evaluate the potential of ViT-based representations alongside prior efforts such as HisToGene [12] and Hist2ST [20].

## 3 EXPERIMENTS AND RESULTS

In this section, we highlight the comprehensive set of experiments undertaken to benchmark different DL architectures for the task of gene expression prediction. We evaluate the performance of the models and provide a novel biological context driven evaluation scheme apart from the traditional quantitative metrics such as SRCC, PCC, MAE, and MSE.

### 3.1 Experiment Setup

Our DL models were trained with a Learning Rate (LR) of 0.0001 using the Adam optimizer for the MSE loss function. The models are trained to predict spatial gene expression at the patch level. Since the encoder backbones were pre-trained on images of size 224 × 224, the input patches were resized to this resolution using standard transformation operations. We applied transfer learning by initializing each model with pretrained weights and finetuning it on the target dataset. All training and inference processes were carried out on a multi-GPU setup consisting of NVIDIA Tesla V100 GPUs (32 GB VRAM) running CUDA version 12.6. This computational setup enabled optimization of deep learning models on high-resolution histopathology images.

### 3.2 Model Performance Comparison

In Table 2 all the models were trained on a patch size 128 × 128, and all models preserved the gene order, which means the predicted gene ranking matches the gene order relationship. The R50 model had the highest SRCC, while EffNet had the highest PCC score. PCC measures the linear relationship between predicted and actual gene expression value. The EffNet model which achieved lowest PCC also had the lowest MSE. R50 provides the lowest MAE. D121, VGG and UNI underperfomed when compared to light-weight EffNet and R50. Moreover, EffNet provided the best trade-off between performance and model size as illustrated in Table 2. It consistently performed better than other models with far fewer parameters.

**Table 2.** Comparison of different model performance on 128 × 128 image patches.

| Metric | EffNet | R50 | D121 | VGG | UNI |
| --- | --- | --- | --- | --- | --- |
| # Parameters (in million) | 5.3 | 25.6 | 8.0 | 138.0 | 300.0 |
| # Layers | 82 | 50 | 121 | 16 | 125 |
| SRCC ↑ | 0.2330 | **0.2338** | 0.2303 | 0.2282 | 0.2246 |
| PCC ↑ | **0.3580** | 0.3497 | 0.3425 | 0.3357 | 0.3268 |
| MAE ↓ | 0.0992 | **0.0969** | 0.1013 | 0.1011 | 0.1022 |
| MSE ↓ | **0.0747** | 0.0756 | 0.0757 | 0.0767 | 0.0783 |

### 3.3 Impact of Patch Resolution on Model Performance

To understand the impact of spatial context on gene expression prediction, we evaluated the model performance across different patch resoltuion. For this analysis, we used EffNet as it outperformed the other encoder models as shown in Table 2. Based on these results, EffNet was a suitable model for systematically exploring the effects of patch size.

We extracted patches of three sizes—16 × 16, 32 × 32, and 128 × 128—centered around each cell, and trained the model separately on each resolution. This experiment was formulated to evaluate whether including a broader tissue structure enhances gene expression prediction. As shown in Table 3, models trained on larger patches, particularly 128 × 128, consistently outperformed smaller ones across the performance metrics.

**Table 3.** Performance of EffNet across varying image patch sizes. The results highlight the impact of spatial context.

| Metric | $128 \times 128$ | $32 \times 32$ | $16 \times 16$ |
| --- | --- | --- | --- |
| SRCC ↑ | **0.2330** | 0.2293 | 0.2235 |
| PCC ↑ | **0.3580** | 0.3510 | 0.3305 |
| MAE ↓ | **0.0992** | 0.0997 | 0.1022 |
| MSE ↓ | **0.0747** | 0.0760 | 0.0782 |

### 3.4 Performance Comparison For Confidently Predicted Genes

Table 4 shows the genes for which each individual model had the highest performance in prediction. We ranked genes according to their performance metric (SRCC and PCC shown in Table 4) across all samples. Based on this, we selected the genes for different top percentiles, i.e. 99%ile, 95%ile, 90%ile and 75%ile percentiles. From Table 4, we observe that EffNet had the highest confidence in PCC. However, for SRCC gene rank-based analysis on 99%ile, R50 worked better, and D121 performed the best on the remaining subsets, achieveing SRCC of 0.6011, 0.5374 and 0.4413 for 95%ile, 90%ile and 75%ile top genes as depicted in Table 4.

**Table 4.** Comparison of model performance for predicted gene expressions across different percentile subsets.

| Metric | %ile | EffNet | R50 | D121 | VGG | UNI |
| --- | --- | --- | --- | --- | --- | --- |
| SRCC ↑ | 99% | 0.7426 | **0.7433** | 0.7408 | 0.7376 | 0.7305 |
|  | 95% | 0.5814 | 0.5909 | **0.6011** | 0.5769 | 0.5651 |
|  | 90% | 0.5178 | 0.5277 | **0.5374** | 0.5124 | 0.5015 |
|  | 75% | 0.4247 | 0.4322 | **0.4413** | 0.4194 | 0.4117 |
| PCC ↑ | 99% | **0.8667** | 0.8642 | 0.8648 | 0.8594 | 0.8506 |
|  | 95% | **0.7765** | 0.7710 | 0.7684 | 0.7607 | 0.7540 |
|  | 90% | **0.7062** | 0.7006 | 0.6987 | 0.6904 | 0.6777 |
|  | 75% | **0.6052** | 0.5982 | 0.5968 | 0.5842 | 0.5717 |

### 3.5 Performance Comparison over Highly Variant Genes

From the dataset, we measured the standard deviation of the individual gene’s expression and ranked them in descending order. The most variant genes are often involved in disease progression and can aid in cell type classification. We compared the performance of the different models for different fractions of the most variant genes. Although the results were close (see Table 5), we observed that predictions from EffNet had the highest PCC whereas predictions from D121 had the best performances for SRCC metrics respectively.

**Table 5.** Model performance on highly variable genes across multiple variance subsets. Top predicted genes metrics reported for SRCC and PCC.

| Metric | Var Top % | EffNet | R50 | D121 | VGG | UNI |
| --- | --- | --- | --- | --- | --- | --- |
| SRCC ↑ | 1% | **0.7244** | 0.7222 | 0.7224 | 0.7179 | 0.7078 |
|  | 5% | 0.5597 | 0.5603 | **0.5622** | 0.5490 | 0.5380 |
|  | 10% | 0.4994 | 0.4988 | **0.5014** | 0.4894 | 0.4783 |
|  | 25% | 0.4060 | 0.4078 | **0.4130** | 0.3978 | 0.3894 |
| PCC ↑ | 1% | **0.8045** | 0.8013 | 0.8006 | 0.7954 | 0.7815 |
|  | 5% | **0.6422** | 0.6345 | 0.6344 | 0.6256 | 0.6075 |
|  | 10% | **0.5929** | 0.5845 | 0.5841 | 0.575 | 0.5556 |
|  | 25% | **0.5117** | 0.5035 | 0.5037 | 0.4927 | 0.4766 |

### 3.6 Predicting Genes Enriched in Biological Pathways

To have biological context as a mode of evaluation, we performed downstream pathway-specific analysis. Broadly, we considered the set of 460 genes for which the ST information were available across the samples as a geneset. The human genome comprises ≈ 20,000 protein-coding genes [21]. These genes were considered as the background genes. We then performed over-representation analysis, often referred to as pathway enrichment analysis [22].

Pathway enrichment analysis involved identifying whether a predefined set of genes (i.e. a known pathway) was overrepresented in a list of genes of interest (460 genes available in our geneset). If a pathway contained a significantly higher number of genes from our geneset of interest than expected by chance, it’s considered enriched. The significance of the enriched pathway was obtained by undertaking a hyper-geometric test. Formally, the goal of hypergeometric test was to compute the probability of observing *k* or more overlapping genes between

- A set of user-selected genes i.e. our 460 ST genes.
- A given biological pathway (e.g., from KEGG [23], Reactome [24], etc.), given the background set of all protein-coding genes.

Mathematically, this can be represented as:

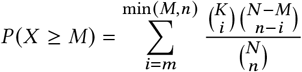

where *N* was the total number of background genes, *M* was the number of genes in a pathway, *n* was the number of genes in our geneset and *m* was number of overlapping genes between a pathway and our geneset. We then performed false discovery rate [25] correction to get q-values for enrichment.

In our work, we used the ConsensusPathDB [26] for the functional and pathway enrichment analysis of our geneset. ConsensusPathDB allowed us to perform overexpression analysis to identify significantly enriched pathways. The advantage of using ConsensusPathDB over a popular tool like DAVID [27] was that it provided the option to search through multiple databases (different types of interactions) to find enriched pathways, unlike DAVID, which only used the KEGG database. Moreover, unlike ingenuity pathway analysis, ConsensusPathDB is a free open source software available for such enrichment an lysis. We included databases such as Biocarta [28], KEGG [23], WikiPathways [29] and Reactome [24], all of which were available in ConsensusPathdb, to identify enriched pathhways. The same pathway can appeared in one of more database. To avoid this we used the ‘clean_pathways’ function from ‘func2vis’ package (v1.0.3) [30] in R (v4.4.1).

Table 6 highlights the top 20 significantly enriched pathways and the performance of the different architectures used to predict gene expression. Table 6 highlight the mean SRCC and PCC respectively for the overlapping genes, between our geneset and the genes in the pathway, based on the correlations between predicted and groundtruth gene expression (for each gene) taking all validation patches into consideration.

**Table 6.** Pathway-wise PCC performance across models. Top 20 significantly enriched pathways are highlighted. Bold values represent the highest PCC for each pathway based on average PCC calculated using all the overlapping genes i.e. #Genes.

| Pathway | p-val | #Genes | EffNet | R50 | D121 | VGG | UNI |
| --- | --- | --- | --- | --- | --- | --- | --- |
| Cytokine-cytokine receptor interaction- Homo sapiens | $10^{-74}$ | 97 | **0.2617** | 0.2557 | 0.2432 | 0.2439 | 0.2351 |
| Viral protein intraxn. cytokine and cytokine receptor- Homo sapiens | $10^{-60}$ | 57 | **0.2847** | 0.2778 | 0.2669 | 0.2657 | 0.2586 |
| Chemokine receptors bind chemokines | $10^{-39}$ | 37 | **0.2859** | 0.2791 | 0.2687 | 0.2675 | 0.2596 |
| Inflammatory bowel disease - Homo sapiens | $10^{-27}$ | 30 | **0.2445** | 0.2366 | 0.2207 | 0.2271 | 0.2197 |
| Selective expression of chemokine receptors during T-cell polarization | $10^{-27}$ | 22 | **0.2463** | 0.2427 | 0.2272 | 0.2323 | 0.2269 |
| Immune System | $10^{-26}$ | 140 | **0.3316** | 0.3233 | 0.3141 | 0.3099 | 0.2963 |
| Hematopoietic cell lineage - Homo sapiens | $10^{-26}$ | 34 | **0.3640** | 0.3528 | 0.3423 | 0.3429 | 0.3292 |
| Allograft Rejection | $10^{-26}$ | 33 | **0.2984** | 0.2879 | 0.2759 | 0.2769 | 0.2662 |
| Th17 cell differentiation - Homo sapiens | $10^{-22}$ | 32 | **0.2942** | 0.2858 | 0.2762 | 0.2780 | 0.2701 |
| JAK-STAT signaling pathway - Homo sapiens | $10^{-22}$ | 38 | **0.2254** | 0.2210 | 0.2058 | 0.2104 | 0.2025 |
| Rheumatoid arthritis - Homo sapiens | $10^{-21}$ | 29 | **0.3059** | 0.2952 | 0.2831 | 0.2817 | 0.2698 |
| IL12-mediated signaling events | $10^{-21}$ | 25 | **0.3319** | 0.3264 | 0.3153 | 0.3149 | 0.3055 |
| Cytokine Signaling in Immune system | $10^{-20}$ | 59 | **0.2727** | 0.2662 | 0.2564 | 0.2543 | 0.2427 |
| Signaling by Interleukins | $10^{-18}$ | 41 | **0.2614** | 0.2567 | 0.2432 | 0.2440 | 0.2309 |
| IL23-mediated signaling events | $10^{-17}$ | 18 | **0.2742** | 0.2695 | 0.2488 | 0.2565 | 0.2447 |
| Development and heterogeneity of the ILC family | $10^{-17}$ | 17 | **0.1798** | 0.1746 | 0.1612 | 0.1615 | 0.1578 |
| G alpha (i) signalling events | $10^{-16}$ | 44 | **0.3063** | 0.2977 | 0.2869 | 0.2841 | 0.2754 |
| IL27-mediated signaling events | $10^{-16}$ | 15 | **0.1842** | 0.1792 | 0.1532 | 0.1668 | 0.1678 |
| Cytokines and Inflammatory Response | $10^{-15}$ | 15 | **0.2371** | 0.2306 | 0.2093 | 0.2156 | 0.2038 |
| Regulatory circuits of the STAT3 signaling pathway | $10^{-15}$ | 22 | **0.3255** | 0.3216 | 0.3184 | 0.3103 | 0.3034 |

Table 6 indicates that several cytokine and chemokine related pathways as well as inflammatory bowel disease and interleukin signaling pathways (*IL12, IL23, IL27*-mediated pathways) are significantly enriched based on our geneset of 460 genes. Using the overlapping set of genes between our geneset and genes which were part of the pathway, we identified a set of biologically relevant genes from that pathway which were part of our predictions. For each suchgene in one such overlapping set, we calculate the PCC between predicted gene expression and groundtruth expression and average the PCC values over all the genes in the overlapping set to get mean PCC prediction for that pathway. From Table 6, we observed that the light-weight EffNet method performs the best for all the significantly enriched pathways, highlighting its supremacy in capturing biologically relevant genes’ expression over other methods like R50, D121, VGG, and UNI.

## 4 DISCUSSION

The rapid advancement of high-resolution ST technologies, such as Xenium, opens new opportunities for modeling spatial gene expression directly from histology images. Our study contributes to this emerging domain by systematically evaluating how model complexity and patch size influence the performance of deep learning models in predicting gene expression from H&E stained images.

One of our key findings is that smaller convolutional models, particularly EffNet, achieved comparable or superior performance to larger models such as R50, D121, VGG, and UNI, especially when trained on larger image patches. Moreover, EffNet initialized with pretrained weights outperforms its counterparts trained from scratch (MSE = 0.0768, SRCC = 0.2278). These results suggest that model depth or parameter count alone does not guarantee improved accuracy. Instead, a more efficient use of spatial context—captured through larger patches that include neighboring cellular information—may be more beneficial for this task. This reinforces the importance of incorporating broader tissue context when modeling spatial gene expression, particularly in heterogeneous tissue samples such as those found in autoimmune diseases.

Despite these promising results, several challenges still need to be addressed. First, co-registration between histopathology and ST data is inherently noisy, due to tissue deformation, staining differences, and spatial resolution mismatches. These factors can introduce uncertainty in model training. Second, ST data remains sparse and noisy, with many transcripts, especially those that are lowly expressed, being underrepresented. This limits the model’s ability to learn accurate mappings for the full gene expression profile. Additionally, cell-type heterogeneity and the absence of explicit segmentation can blur morphological signals, making it harder to associate image features with specific molecular signatures. Importantly, some genes are regulated by non-morphological factors such as the local microenvironment, immune signaling, or temporal dynamics, which are not captured in static H&E images. This presents a fundamental limitation for purely image-based prediction.

Future extensions of the work can handle some of these biological and technical limitations. In particular, we plan to include the interpretability methods, such as attention map visualization and gene saliency analysis, to better understand the morphological cues driving model predictions. Expanding the evaluation beyond the AMLC dataset to other tissue types and spatial transcriptomics platforms, such as 10x Genomics Visium, will also be critical for measuring the generalizability of our findings. Additional directions include the integration of single-cell RNA-seq data, multi-scale image features, and / or graph-based modeling of cellular neighborhoods. We could also look into using ordinal regression frameworks from medical imaging [31] could be considered for task-specific gene ranking problems that may emerge in ST tasks, as these models are good to capture ordered relationships between prediction. Together, these efforts can provide a more comprehensive framework for spatial gene expression prediction. Ultimately, the ability to recover spatially resolved gene expression from standard histopathology images has broad implications for diagnostics, biomarker discovery, and the virtual extension of ST technologies.

Overall, our findings provide actionable guidance for designing lightweight yet effective models for spatial gene expression prediction. This can support broader adoption of virtual ST technologies in research and clinical workflows, enabling cost-effective molecular profiling from standard histopathology images.

## Notes

### Competing Interest Statement

The authors have declared no competing interest.

### Summary of Updates

I think we accidentally missed Dr. Raghvendra's name in the PDF. He was added in the system, though. Have submitted the revised PDF.

